# NAIP–NLRC4-deficient mice are susceptible to shigellosis

**DOI:** 10.1101/2020.05.16.099929

**Authors:** Patrick S. Mitchell, Justin L. Roncaioli, Elizabeth A. Turcotte, Lisa Goers, Roberto A. Chavez, Angus Y. Lee, Cammie F. Lesser, Isabella Rauch, Russell E. Vance

## Abstract

Bacteria of the genus *Shigella* cause shigellosis, a severe gastrointestinal disease that is a major cause of diarrhea-associated mortality in humans. Shigellosis develops upon oral ingestion of as few as 100 bacteria, but million-fold higher doses fail to cause disease in mice. The lack of a physiologically relevant mouse model of shigellosis has impeded our understanding of this important human disease, but why mice are resistant is unknown. Here we show that in human cells, but not in mice, *Shigella* evades detection by the NAIP–NLRC4 inflammasome, an immune sensor present in intestinal epithelial cells (IECs). We find that NAIP–NLRC4-deficient mice are highly susceptible to oral *Shigella* infection and recapitulate the clinical features of human shigellosis, including bacterial replication in IECs and neutrophilic inflammation of the colon. Confirming a role for bacterial replication in IECs in our new model, a *Shigella* mutant lacking IcsA, a factor required for cell-to-cell spread among IECs, is attenuated in otherwise susceptible NAIP–NLRC4-deficient mice. Although inflammasome-mediated cell death is widely held to promote *Shigella* infection and pathogenesis, we instead demonstrate that IEC-specific NAIP–NLRC4-induced cell death is sufficient to protect the host from shigellosis. Thus, NAIP–NLRC4-deficient mice are a physiologically relevant and experimentally tractable model for shigellosis. More broadly, our results suggest that the lack of an inflammasome response in IECs may help explain the extreme susceptibility of humans to shigellosis.

## Introduction

*Shigella* is a genus of Gram-negative enterobacteriaceae that causes ∼269 million infections and ∼200,000 deaths annually, a quarter of which are of children under the age of five (Khalil et al., 2018). Disease symptoms include fever, abdominal cramping, and inflammatory diarrhea characterized by the presence of neutrophils and, in severe cases, blood (Kotloff et al., 2018). There is no approved vaccine for *Shigella* and antibiotic resistance continues to rise (Ranjbar and Farahani, 2019). *Shigella* pathogenesis is believed to be driven by bacterial invasion, replication, and spread within colonic IECs. *Shigella* virulence requires a plasmid-encoded type III secretion system (T3SS) that injects ∼30 effectors into host cells (Schnupf and Sansonetti, 2019; Schroeder and Hilbi, 2008). The virulence plasmid also encodes IcsA, a bacterial surface protein that nucleates host actin at the bacterial pole to propel the pathogen through the host cell cytosol and into adjacent epithelial cells (Bernardini et al., 1989; Goldberg and Theriot, 1995).

A major impediment to studying *Shigella* is the lack of experimentally tractable *in vivo* models that accurately recapitulate human disease after oral inoculation. Although the infectious dose for humans is as low as 10-100 bacteria (DuPont et al., 1969; DuPont et al., 1989), mice are resistant to high doses of oral *Shigella* challenge (Freter, 1956; McGuire and Floyd, 1958). Rabbits, guinea pigs, zebrafish, piglets, and macaques have been used as models (Islam et al., 2014; Jeong et al., 2010; Mostowy et al., 2013; Ranallo et al., 2014; Shim et al., 2007; West et al., 2005; Yum and Agaisse, 2019; Yum et al., 2019) but the cost and/or limited tools in these systems impair detailed studies of pathogenesis. Oral streptomycin administration and other treatments facilitate *Shigella* colonization of the mouse intestinal lumen by ablating the natural colonization resistance provided by the microbiome (Freter, 1956; Martino et al., 2005; Medeiros et al., 2019). However, antibiotic-treated mice do not present with key hallmarks of human disease, likely due to the failure of *Shigella* to invade and/or establish a replicative niche within the mouse intestinal epithelium.

Inflammasomes are cytosolic multi-protein complexes that initiate innate immune responses upon pathogen detection or cellular stress (Lamkanfi and Dixit, 2014; Rathinam and Fitzgerald, 2016). The NAIP–NLRC4 inflammasome is activated when bacterial proteins, such as flagellin or the rod and needle proteins of the T3SS apparatus, are bound by NAIP family members. Importantly, the *Shigella* T3SS inner rod (MxiI) and needle (MxiH) proteins are both potent agonists of human and mouse NAIPs (Reyes Ruiz et al., 2017; Yang et al., 2013). Activated NAIPs then co-assemble with NLRC4 to recruit and activate the Caspase-1 (CASP1) protease (Vance, 2015; Zhao and Shao, 2015). CASP1 then cleaves and activates the pro-inflammatory cytokines IL-1β and IL-18 and the pore-forming protein Gasdermin-D (Kayagaki et al., 2015; Shi et al., 2015), initiating a lytic form of cell death called pyroptosis. We and others recently demonstrated that activation of NAIP–NLRC4 in IECs further mediates the cell-intrinsic expulsion of infected epithelial cells from the intestinal monolayer (Rauch et al., 2017; Sellin et al., 2014). In the context of *Shigella* infection, it is generally accepted that inflammasome-mediated pyroptosis of infected macrophages promotes pathogenesis by initiating inflammation, and by releasing bacteria from macrophages, allowing the bacteria to invade the basolateral side of intestinal epithelial cells (Ashida et al., 2014; Lamkanfi and Dixit, 2010; Schnupf and Sansonetti, 2019). However, it has not been possible to test the role of inflammasomes in the intestine after oral *Shigella* infection due to the lack of a genetically tractable model. Here we develop the first oral infection mouse model for *Shigella* infection and demonstrate a specific host-protective function for inflammasomes in intestinal epithelial cells.

## Results

### *Shigella* suppresses the human NAIP–NLRC4 inflammasome

The *Shigella* T3SS effector OspC3 inhibits cytosolic LPS sensing by the human Caspase-4 (CASP4) inflammasome, but does not bind to the mouse ortholog, Caspase-11 (CASP11) (Kobayashi et al., 2013). We reasoned that inflammasome inhibition may be a general strategy used by *Shigella* to establish infection, and that such inhibition might occur in a host-specific manner. To test this hypothesis, we compared inflammasome-dependent cell death following *Shigella* infection of mouse C57BL/6 (B6) bone marrow-derived macrophages (BMMs) and human PMA-differentiated THP1 cells. Infection with the wild-type (WT) *Shigella flexneri* strain 2457T but not the avirulent BS103 strain (which lacks the virulence plasmid) resulted in CASP1-dependent cell death in both mouse (Sandstrom et al., 2019) and human cells (**Figure 1A,B**). Cell death was negligible in *Shigella*-infected mouse *Nlrc4*^*–/–*^ BMMs (**Figure 1A**). In contrast, *Shigella* infection induced similar levels of cell death in WT and *NLRC4*^*–/–*^ THP1 cells, indicating that NLRC4 is not a major contributor to *Shigella*-induced CASP1 activation in human cells (**Figure 1B**).

**Figure 1.**
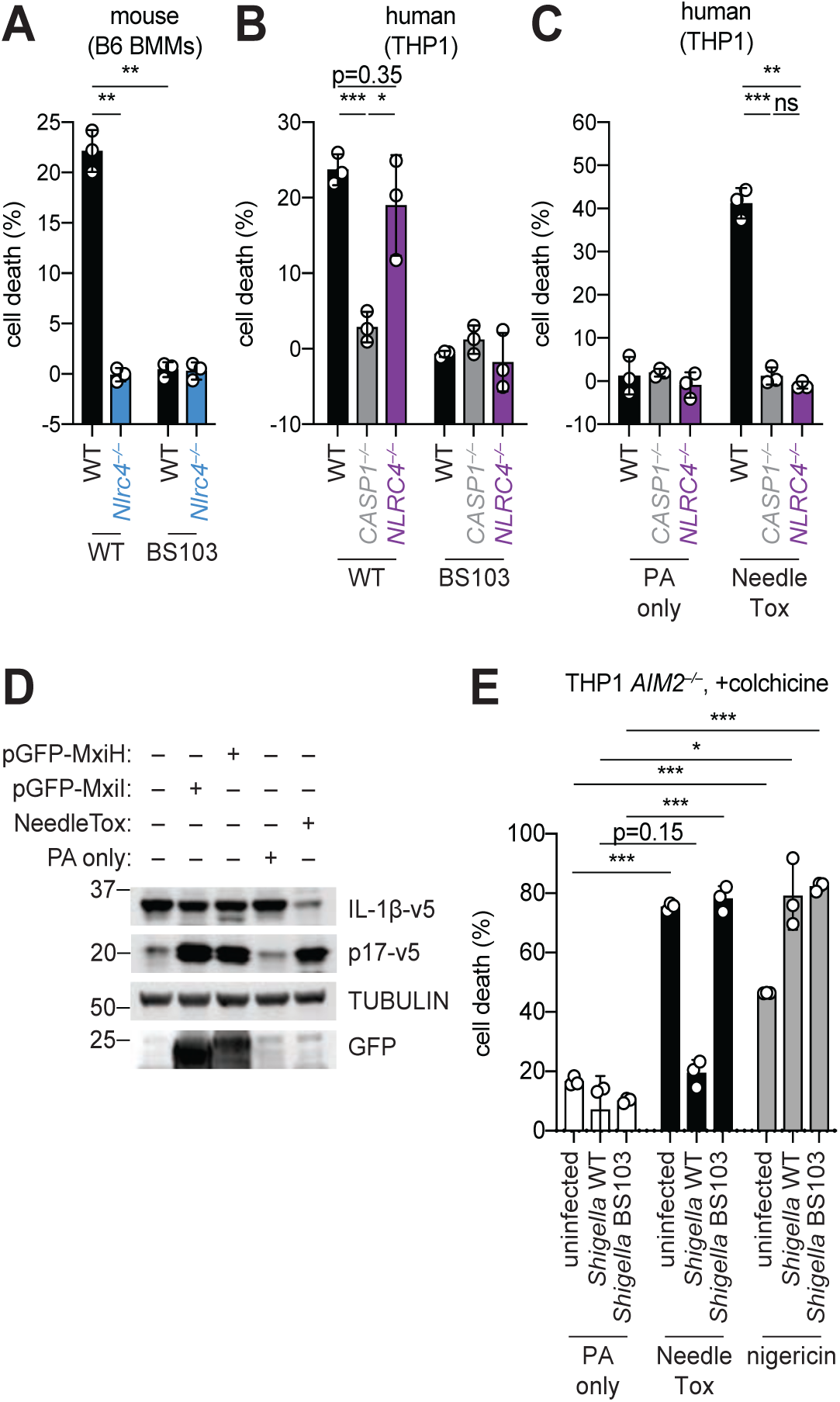
*Shigella* infection suppresses the NAIP–NLRC4 inflammasome. (**A**) *Shigella* infection (MOI 10) of C57BL/6 WT or *Nlrc4*^*–/–*^ bone marrow derived macrophages (BMMs). Cell death was measured 30 minutes post-infection (after spinfection, invasion, and washes) by propidium iodide uptake and reported as percent death relative to 100% killing by treatment with Triton X-100. (**B**) Cell death of *Shigella* infected THP1 WT, *CASP1*^*–/–*^ or *NLRC4*^*–/–*^ cells as in (**A**). Cell death was measured 30 minutes post-infection. (**C**) Cell death of THP1 WT, *CASP1*^*–/–*^ or *NLRC4*^*–/–*^ cells treated with 10µg/mL PA alone or in combination with 10µg/mL LFn-MxiH (“NeedleTox”). Cell death was measured 4 hours post-challenge. (**D**) Human NAIP–NLRC4 inflammasome reconstitution in 293T cells. Inflammasome activation was measured by CASP1-dependent processing of pro-IL-1β to p17 by co-transfection of an empty vector, pGFP-MxiH or pGFP-MxiI, or by treatment with 10µg/mL PA alone or in combination with 10µg/mL LFn-MxiH. (**E**) Colchicine-treated *AIM2*^*–/–*^ THP1 cells were either left uninfected or infected for 1 hour (after spinfection, invasion, and washes) with WT or BS103 *Shigella* (MOI 10), and then treated with 10µg/mL PA alone, PA + 1.0µg/mL LFn-MxiH (“NeedleTox”), or 10µM nigericin. Cell death was measured by PI staining and is reported as cell death relative to TX-100-treated controls per infection type. Data are representative of at least three independent experiments. Mean ± SD is shown in (**A–C,E**), unpaired t-test with Welch’s correction: \**P* < 0.01, \*\**P* < 0.001, ***P < 0.0001.

To confirm prior reports (Yang et al., 2013) that the *Shigella* T3SS MxiH (needle) protein is a potent agonist of the human NAIP–NLRC4 inflammasome, and that THP1 cells express a functional NAIP–NLRC4 inflammasome (Kortmann et al., 2015; Reyes Ruiz et al., 2017), we produced recombinant LFn-MxiH for cytosolic delivery to cells via the protective antigen (PA) channel, as previously described (Rauch et al., 2017; Rauch et al., 2016; von Moltke et al., 2012). Treatment of THP1 cells with LFn-MxiH+PA (“NeedleTox”), but not PA alone, induced a CASP1- and NLRC4-dependent cell death (**Figure 1C**). We also confirmed that *Shigella* MxiH and MxiI (rod) activate human NAIP–NLRC4 in a reconstituted inflammasome assay (Reyes Ruiz et al., 2017; Tenthorey et al., 2014) (**Figure 1D**). These results confirm that the *Shigella* rod and needle proteins are capable of activating the human NAIP–NLRC4 inflammasome. Moreover, since we observe robust activation of NAIP–NLRC4 in mouse cells (**Figure 1A**), our data indicate that the *Shigella* T3SS needle and rod proteins are delivered to the cytosol during infection. However, the presence of other inflammasomes in THP1 cells obscures our ability to determine if *Shigella* activates human NLRC4.

To eliminate cell death induced by the AIM2 and PYRIN inflammasomes, we used THP1 *AIM2*^*–/–*^ cells treated with colchicine, an inhibitor of PYRIN (Gao et al., 2016). Interestingly, WT *Shigella* infection did not induce pyroptosis of colchicine-treated *AIM2*–/– THP1 cells (**Figure 1E**). Thus, although human THP1 cells express functional NAIP–NLRC4, and *Shigella* rod and needle proteins can activate NAIP–NLRC4, we nevertheless observe no such activation during *Shigella* infection. We therefore hypothesized that *Shigella* might antagonize the human NAIP-NLRC4 inflammasome. To test this hypothesis, *AIM2*^*–/–*^ colchicine-treated THP1 cells were either uninfected or infected with the *Shigella* WT or BS103 strains for one hour, and then treated with NeedleTox to induce NAIP-NLRC4-dependent cell death. Interestingly, NeedleTox-induced pyroptosis was significantly reduced in cells previously infected with WT but not avirulent BS103 *Shigella* (**Figure 1E**). As a control, treatment with nigericin, an agonist of the NLRP3 inflammasome, induced cell death similarly in cells that were uninfected or infected with either WT or BS103 *Shigella* strains. Thus, WT *Shigella* infection appears to specifically suppress activation of the human but not the mouse NAIP– NLRC4 inflammasome. Future studies will be required to address the mechanism of *Shigella* antagonism of human NAIP–NLRC4, as well as the mechanism of *Shigella* activation of AIM2 and/or PYRIN.

### B6.*Naip*-deficient mice are susceptible to shigellosis

The mouse NAIP–NLRC4 and CASP11 inflammasomes protect the intestinal epithelium from *Salmonella* (Rauch et al., 2017; Sellin et al., 2014). Thus, the above experiments led us to hypothesize that the failure of *Shigella* to antagonize mouse inflammasomes might explain the inborne resistance of mice versus humans to *Shigella* infection. A prediction from this hypothesis is that mice deficient in inflammasomes might be susceptibility to oral *Shigella* challenge.

To test this, we first pretreated B6 WT mice orally with streptomycin antibiotic. Consistent with prior studies (Freter, 1956; Martino et al., 2005; Medeiros et al., 2019), we found that antibiotic pre-treatment followed by oral infection allows for robust *Shigella* colonization of the intestinal lumen and feces compared to water-only controls (**Figure S1**). However, these high lumenal bacterial loads (>10^8^ CFU/g feces) do not cause overt disease (**Figure 2** and **Figure S2**). To determine if inflammasomes contribute to the resistance of mice to *Shigella*, we orally challenged WT and *Casp1/11*^*–/–*^ mice with 5×10^7^ CFU of *Shigella*. CASP1 has been previously reported to drive acute inflammation and *Shigella* clearance during mouse lung infection via the processing and release of IL-1β and IL-18 (Sansonetti et al., 2000). Mice that lack CASP1 and/or CASP11 are also more susceptible to oral infection with *Salmonella* Typhimurium (Crowley et al., 2020). In contrast, B6 WT and *Casp1/11*^*–/–*^ mice were similarly resistant to *Shigella* infection, showing no signs of intestinal inflammation or disease (**Figure S2A–C**). Thus, neither of the primary caspases associated with the canonical or non-canonical inflammasome are essential for resistance to *Shigella* in the mouse intestine.

**Figure 2.**
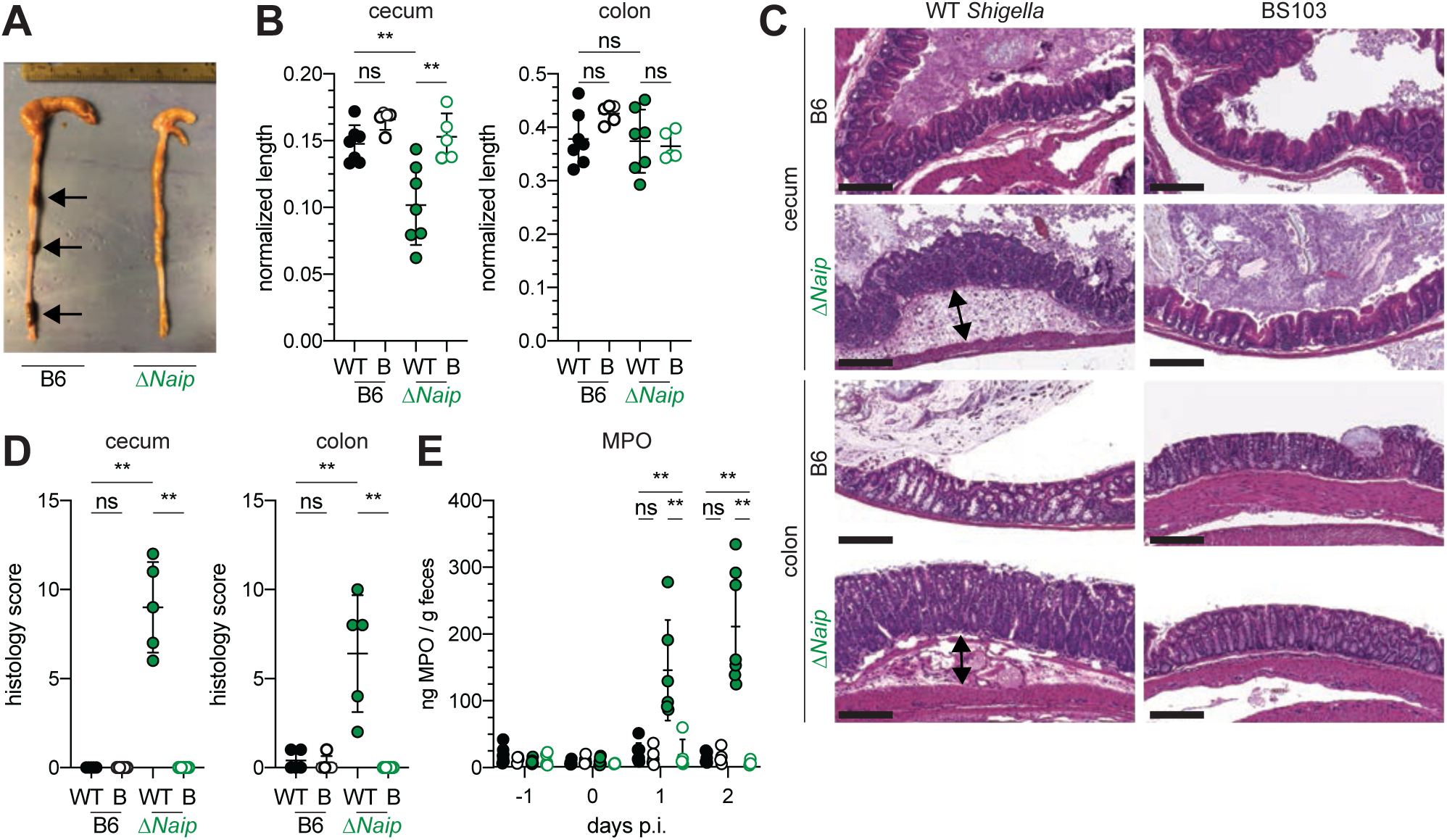
*Shigella*-infected B6.Δ*Naip* mice exhibit intestinal inflammation. (**A–E**) B6.WT and B6.Δ*Naip* (green) mice (lacking expression of all *Naip* genes) treated orally with 25mg streptomycin sulfate were orally challenged the next day with 5×10^7^ CFU of WT or BS103 (non-invasive) *Shigella*. Endpoint harvests were performed at 48 hours post-infection (p.i.). (**A**) Representative images of the cecum and colon dissected from B6.WT and B6. Δ*Naip* mice. Note cecum tissue thickening (size reduction), macroscopic edema, and loose stool (absence of arrows). (**B**) Quantification of cecum and colon lengths. Values were normalized to mouse weight prior to infection; cecum length (cm) / mouse weight (g). WT, wild-type *Shigella* (filled symbols); B, BS103 (open symbols). (**C**) Representative images of H&E stained cecum and colon tissue from infected mice. Scale bar, 200 µm. (**D**) Blinded quantification of histology score (cumulative) for tissues in (**C**). Edema, hyperplasia, inflammatory infiltrate, and epithelial cell death were scored from 0-4. The final score is the sum of individual scores from each category. (**E**) MPO levels measured by ELISA from feces of B6.WT and B6.Δ*Naip* mice collected -1 through 2 days p.i. (**B, D, E**) Each symbol represents one mouse. Filled symbols, WT *Shigella*; open symbols, BS103. Data are representative of two independent experiments. Mean ± SD is shown in (**B,D,E**), Mann-Whitney test, \**P* < 0.05, \*\**P* < 0.01, \*\*\**P* < 0.001.

The NAIP–NLRC4 inflammasome can recruit Caspase-8 (CASP8) in the absence of CASP1, an event which leads to non-lytic cell death and delayed IEC expulsion (Rauch et al., 2017). We reasoned that this compensatory capacity of CASP8 may account for the resistance of *Casp1/11*^*–/–*^ mice to *Shigella* infection. Thus, to directly test if the mouse NAIP–NLRC4 inflammasome mediates resistance to shigellosis, we orally infected streptomycin-pretreated B6 WT and Δ*Naip* mice with 5×10^7^ CFU of *Shigella*. Δ*Naip* mice (also called *Naip1-6*Δ/Δ mice (Rauch et al., 2016)) harbor a large chromosomal deletion that eliminates expression of all mouse *Naip* genes. Remarkably, *Shigella*-infected Δ*Naip* but not WT mice exhibited clear signs of disease (**Figure 2**). At two days post-challenge, *ΔNaip* mice had altered stool consistency (**Figure 2A**), cecum shrinkage, and thickening of the cecum and colon tissue (**Figure 2A,B**). Histological analysis of primary sites of infection (cecum, colon) revealed edema, epithelial hyperplasia, epithelial sloughing, and inflammatory infiltrate (predominantly neutrophils and mononuclear cells in the submucosa and mucosa) exclusively in Δ*Naip* mice (**Figure 2C,D**). In contrast, we did not observe any indicators of inflammation in Δ*Naip* mice infected with the avirulent BS103 strain (**Figure 2C,D**).

A defining feature of human shigellosis is the presence of neutrophils in patient stools (Raqib et al., 2000). The levels of myeloperoxidase (MPO, a neutrophil marker) were low or undetectable in the feces of mice following antibiotic treatment (**Figure 2E**), indicating that microbiota disruption did not itself promote neutrophilic inflammation. Following *Shigella* infection, however, fecal MPO from Δ*Naip* mice dramatically increased (**Figure 2E**). In contrast, MPO levels remained low in both B6 WT and *Casp1/11*^*–/–*^ mice (**Figure S2D**) or *ΔNaip* mice infected with the avirulent BS103 strain (**Figure 2E**). These results indicate that NAIP–NLRC4-deficient mice experience robust neutrophilic infiltrate consistent with human shigellosis.

### B6.*Nlrc4*^−/−^ mice are susceptible to shigellosis

To confirm that the NAIP–NLRC4 inflammasome confers resistance to *Shigella*, to account for potential microbiota-associated phenotypes, and to further characterize the disease phenotype, we next infected streptomycin-pretreated B6.*Nlrc4*^+/–^ and B6.*Nlrc4*^−/−^ littermates, as well as B6 WT mice that had been co-housed for three weeks prior to inoculation (*Nlrc4*^+*/–*^ and B6 WT mice are hereby referred to collectively as *Nlrc4*^+^). Consistent with our prior results in Δ*Naip* mice, we observed thickening of the intestinal mucosa (**Figure 3A**), cecum shrinkage (**Figure 3A,B**), increased fecal MPO levels (**Figure 3C**), and acute weight loss (**Figure 3D**) in *Shigella*-infected B6.*Nlrc4*^−/−^ mice but not B6.*Nlrc4*^+^ littermates or co-housed mice. B6.*Nlrc4*^−/−^ mice also had diarrhea, which was apparent by visual inspection of lumenal contents and measured by the wet-to-dry ratio of fecal pellets (**Figure 3E**). Thus, B6.*Nlrc4*^−/−^ mice phenocopy the disease susceptibility of B6.Δ*Naip* mice, and strongly suggest that the NAIP–NLRC4 inflammasome mediates the resistance of mice to *Shigella* infection.

**Figure 3.**
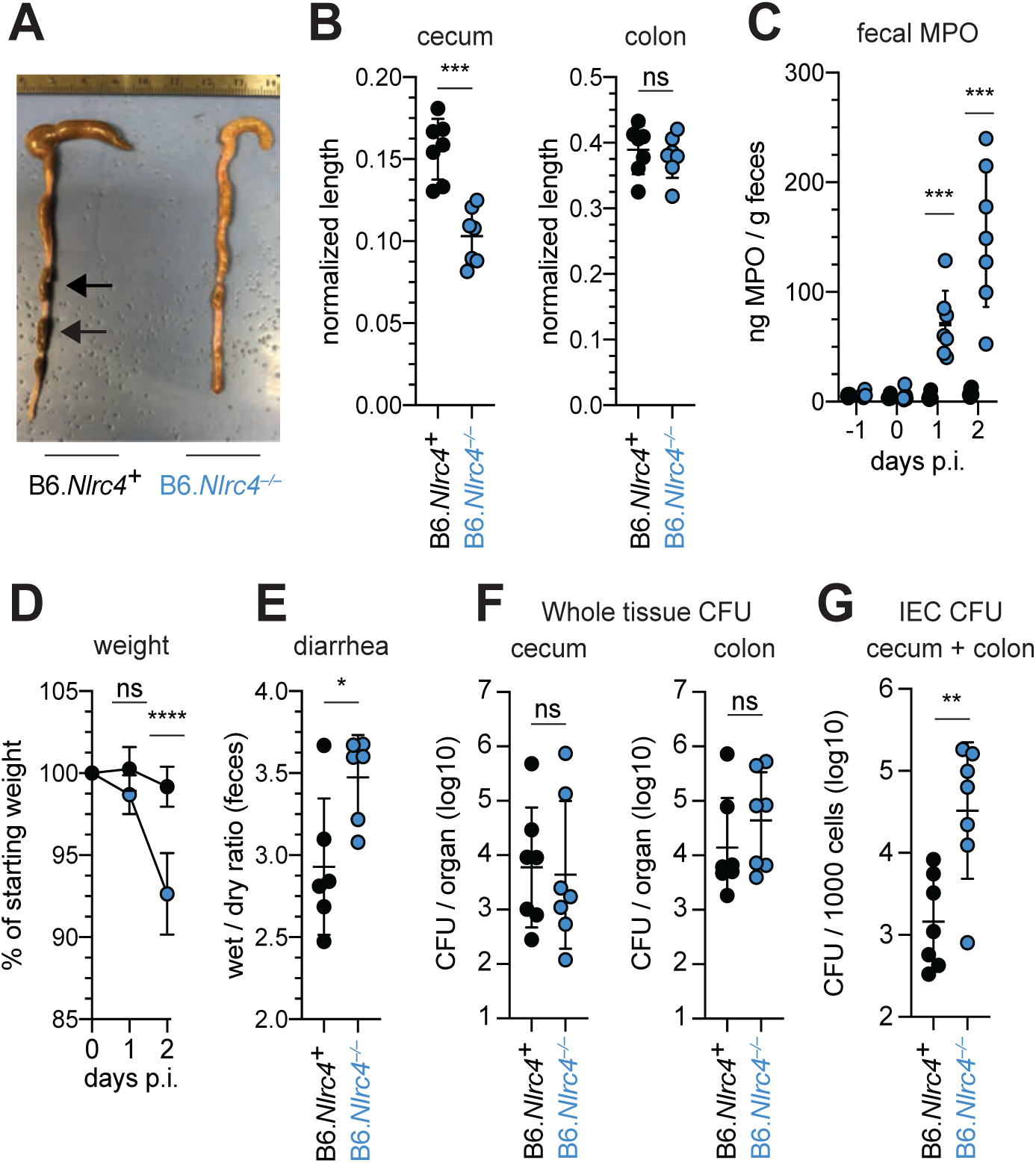
*Shigella*-infected B6.*Nlrc4*^*–/–*^ mice exhibit intestinal inflammation and bacterial colonization of IECs. (**A–E**) B6.*Nlrc4*^+*/–*^ and B6.*Nlrc4*^*–/–*^ littermates were cohoused with B6.WT mice for a minimum of three weeks. Mice were infected as described for Figure 2. Endpoint harvests were performed 48 hours post-infection (p.i.). B6.*Nlrc4*^+*/–*^ and B6.WT mice are collectively referred to as B6.*Nlrc4*^+^. (**A**) Representative images of the cecum and colon dissected from B6.*Nlrc4*^+^ and B6.*Nlrc4*^*–/–*^ mice. Note the cecum tissue thickening (size reduction), macroscopic edema, and loose stool (absence of arrows). (**B**) Quantification of cecum and colon lengths. Values were normalized to mouse weight prior to infection; cecum length (cm) / mouse weight (g). (**C**) MPO levels measured by ELISA from feces of B6.*Nlrc4*^+^ and B6.*Nlrc4*^*–/–*^ mice collected -1 through 2 days p.i. (**D**) Mouse weights from 0 through 2 days p.i. Each symbol represents the mean for all mice of the indicated condition. (**E**) Quantification of feces weights before and after dehydration at 2 days p.i. A larger ratio indicates diarrhea. (**F**) CFU determination from gentamicin-treated whole tissue homogenates from the cecum or colon of infected mice. (**G**) CFU determination from the IEC enriched fraction of gentamicin-treated cecum and colon tissue (combined). (**B,C,E–G**) Each symbol represents one mouse. Data are representative of three independent experiments. Mean ± SD is shown in (**B,D,E**), Mann-Whitney test, \**P* < 0.05, \*\**P* < 0.01, \*\*\**P* < 0.001.

Surprisingly, despite the clear differences in disease between *Shigella* infected B6.*Nlrc4*^+^ and B6.*Nlrc4*^*–/–*^ mice, we found no significant difference in the bacterial burdens of cecum or colon tissue (**Figure 3F**). To more directly measure the intracellular colonization of IECs, the primary replicative niche for *Shigella*, we enriched IECs from the ceca and colons of infected B6.*Nlrc4*^+^ and B6.*Nlrc4*^−/−^ mice (see Methods). We found an ∼20-fold difference in colonization in enriched IECs between B6.*Nlrc4*^+^ and B6.*Nlrc4*^−/−^ mice (**Figure 3G**), indicating that disease in our model correlates with invasion of and replication in IECs. Importantly, we observed no CFU differences in feces at the time of harvest (**Figure S3A**), excluding the possibility that differences in IEC CFU were caused by differences in lumenal *Shigella* density. These data suggest that *Shigella* colonizes the intestinal tissue during infection of either genotype but can only invade the epithelium and provoke disease in NAIP–NLRC4-deficient mice.

### *Shigella* causes bloody diarrheal disease in 129.*Nlrc4*^−/−^ mice

To determine whether the role of NAIP–NLRC4 in mediating protection against *Shigella* is robust across diverse mouse strains, we generated *Nlrc4*^−/−^ mice on the 129S1 genetic background. 129.*Nlrc4*^−/−^ mice have a 10bp deletion in exon 5 of the *Nlrc4* coding sequence, resulting in loss of NLRC4 function (**Figure S4**). Importantly, 129S1 mice are naturally deficient in CASP11, which responds to cytosolic LPS, and thus 129.*Nlrc4*^−/−^ mice lack functional NAIP–NLRC4 and CASP11 signaling.

Similar to the B6 genetic background, antibiotic-pretreated 129.*Nlrc4*^−/−^ but not 129.*Nlrc4*^+/–^ littermates challenged with WT (or BS103) *Shigella* exhibited severe signs of shigellosis, including pronounced edema, epithelial cell hyperplasia, and disruption of the columnar epithelium of infected tissues (**Figure 4A,B**). 129.*Nlrc4*^−/−^ mice also exhibited dramatic cecum shrinkage and diarrhea (**Figure 4C,D,K**), lost between eight and 18 percent of their starting weight within two days of infection (**Figure 4E**), and exhibited a massive increase in fecal MPO following infection (**Figure 4F**). We found no significant difference in the bacterial colonization of the whole cecum and colon tissue between 129.*Nlrc4*^+/–^ and 129.*Nlrc4*^−/−^ mice (**Figure 4G**). However, IECs enriched from infected 129.*Nlrc4*^−/−^ mice again exhibited an ∼20-fold higher bacterial burden than IECs enriched from 129.*Nlrc4*^+/–^ mice (**Figure 4H**), despite similar levels of lumenal colonization (**Figure S3B**).

**Figure 4.**
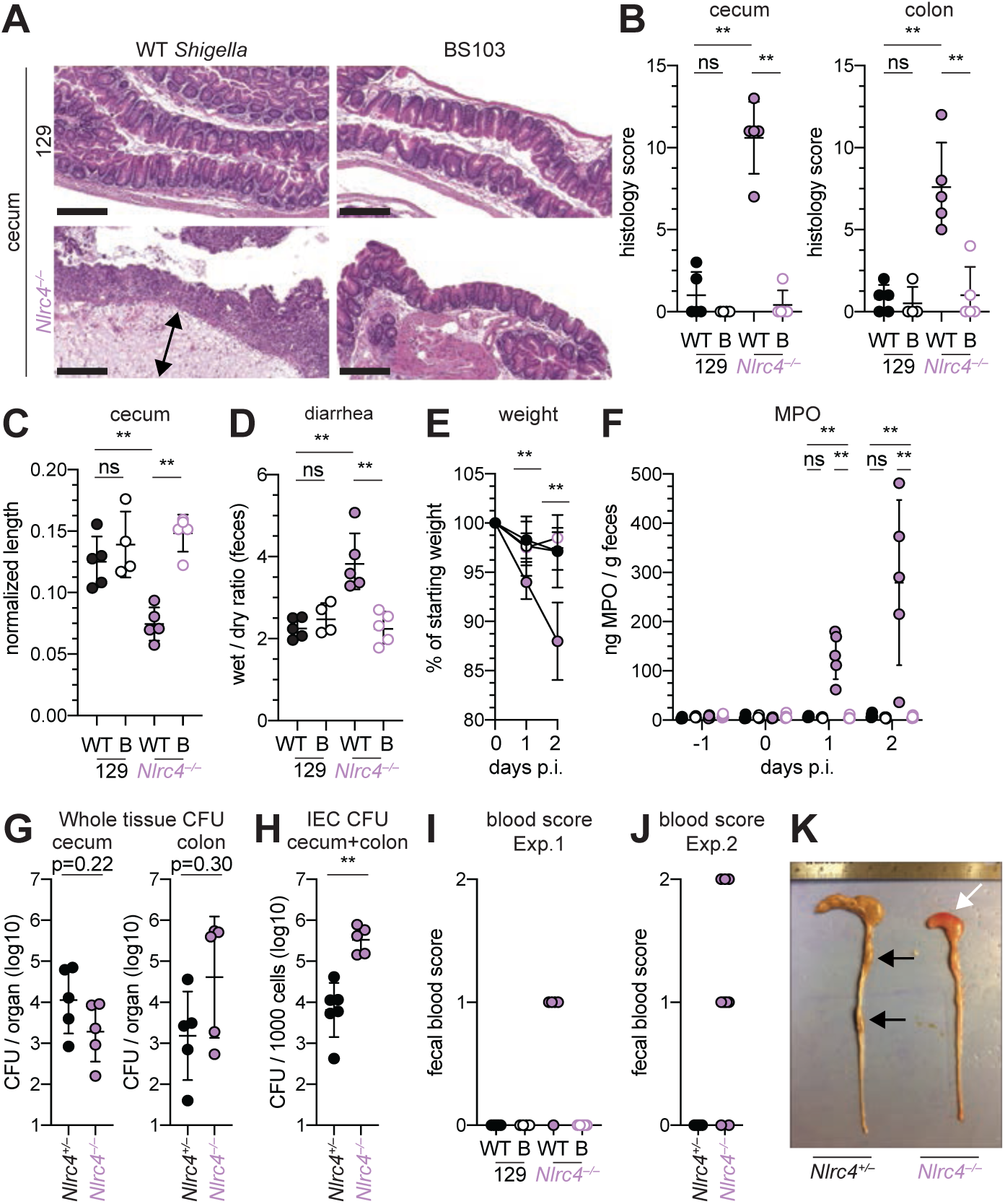
*Shigella*-infected 129.*Nlrc4*^*–/–*^ mice exhibit hallmarks of severe human shigellosis. (**A–H**) 129.*Nlrc4*^+*/–*^ and 129.*Nlrc4*^*–/–*^ littermates were infected as described for **Figure 2**. Endpoint harvests were performed at 48 hours post-infection (p.i.). (**A**) Representative images of H&E stained cecum and colon tissue from infected mice. Scale bar, 200µm. (**B**) Blinded quantification of histology score (cumulative) for tissues in (**A**). Edema, hyperplasia, inflammatory infiltrate, and epithelial cell death were scored from 0-4. The final score is the sum of individual scores from each category. (**C**) Quantification of cecum and colon lengths. Values were normalized to mouse weight prior to infection; cecum length (cm) / mouse weight (g). (**D**) Quantification of feces weights before and after dehydration at two days p.i. A larger ratio indicates diarrhea. (**E**) Mouse weights at 0 through 2 days p.i. Each symbol represents the mean for all mice of the indicated condition. Statistics refer to both WT *Shigella*-infected 129.*Nlrc4*^+*/–*^ and 129.*Nlrc4*^*–/–*^ mice and WT versus BS103 *Shigella*-infected 129.*Nlrc4*^*–/–*^ mice at both 1 and 2 days p.i. All other comparisons were non-significant. (**F**) MPO levels measured by ELISA from feces of 129.*Nlrc4*^+*/–*^ and 129.*Nlrc4*^*–/–*^ mice collected -1 through 2 days p.i. (**G**) CFU determination from gentamicin-treated whole tissue homogenates from the cecum or colon (**H**) CFU determination from the IEC enriched fraction of gentamicin-treated cecum and colon tissue (combined). (**I,J**) Fecal blood scores from feces at two days p.i. 1 = occult blood, 2 = macroscopic blood. (**I**) and (**J**) show scores from two representative experiments. (**K**) Representative images of the cecum and colon dissected from 129.*Nlrc4*^+*/–*^ and 129.*Nlrc4*^*–/–*^ mice. Note the cecum tissue thickening (size reduction), macroscopic edema, and loose stool (absence of arrows), and vascular lesions and bleeding. (**B–D,F**–**J**) Each symbol represents one mouse. Filled symbols, WT *Shigella*; open symbols, BS103. Data are representative of three independent experiments. Mean ± SD is shown in (**B,D,E**), Mann-Whitney test, \**P* < 0.05, \*\**P* < 0.01, \*\*\**P* < 0.001.

A hallmark of severe human shigellosis (dysentery) is the presence of blood in patient stools — a phenotype we did not observe in NAIP–NLRC4-deficient mice on the B6 background. We tested 129.*Nlrc4*^−/−^ mouse stools for the presence of occult blood and found that 4/5 mice infected with WT *Shigella* had occult blood in their feces (**Figure 4I**). In a subsequent infection, 80% (8/10) of 129.*Nlrc4*^*–/–*^ mice had bloody stool (occult blood only, n=5; macroscopically visible blood, n=3) (**Figure 4J**). In mice with visible blood, we often observed ruptured blood vessels in the cecum or colon (**Figure 4K**).

### Epithelial NLRC4 is sufficient to protect mice from shigellosis

Given the difference in *Shigella* colonization of IECs between WT and NAIP–NLRC4-deficient mice, we next sought to determine if IEC-specific expression of the NAIP–NLRC4 inflammasome is sufficient to protect mice from *Shigella* infection. We thus infected B6 mice that selectively express NLRC4 in IECs. These mice encode a Cre-inducible *Nlrc4* gene on an otherwise *Nlrc4*^*–/–*^ background and are referred to as iNLRC4 mice (**Figure 5A**) (Rauch et al., 2017). Crosses of iNLRC4 and *Vil1-Cre* mice generated animals with selective expression of NLRC4 in Villin^+^ IECs. *Shigella* infected *Vil1-Cre*^+^ iNLRC4 mice, but not *Cre*^*–*^ littermate controls, were protected from intestinal inflammation to a similar extent as co-housed *Nlrc4*^+*/–*^ mice (**Figure 5B–F**). Thus, NLRC4 expression in IECs is sufficient to prevent shigellosis.

**Figure 5.**
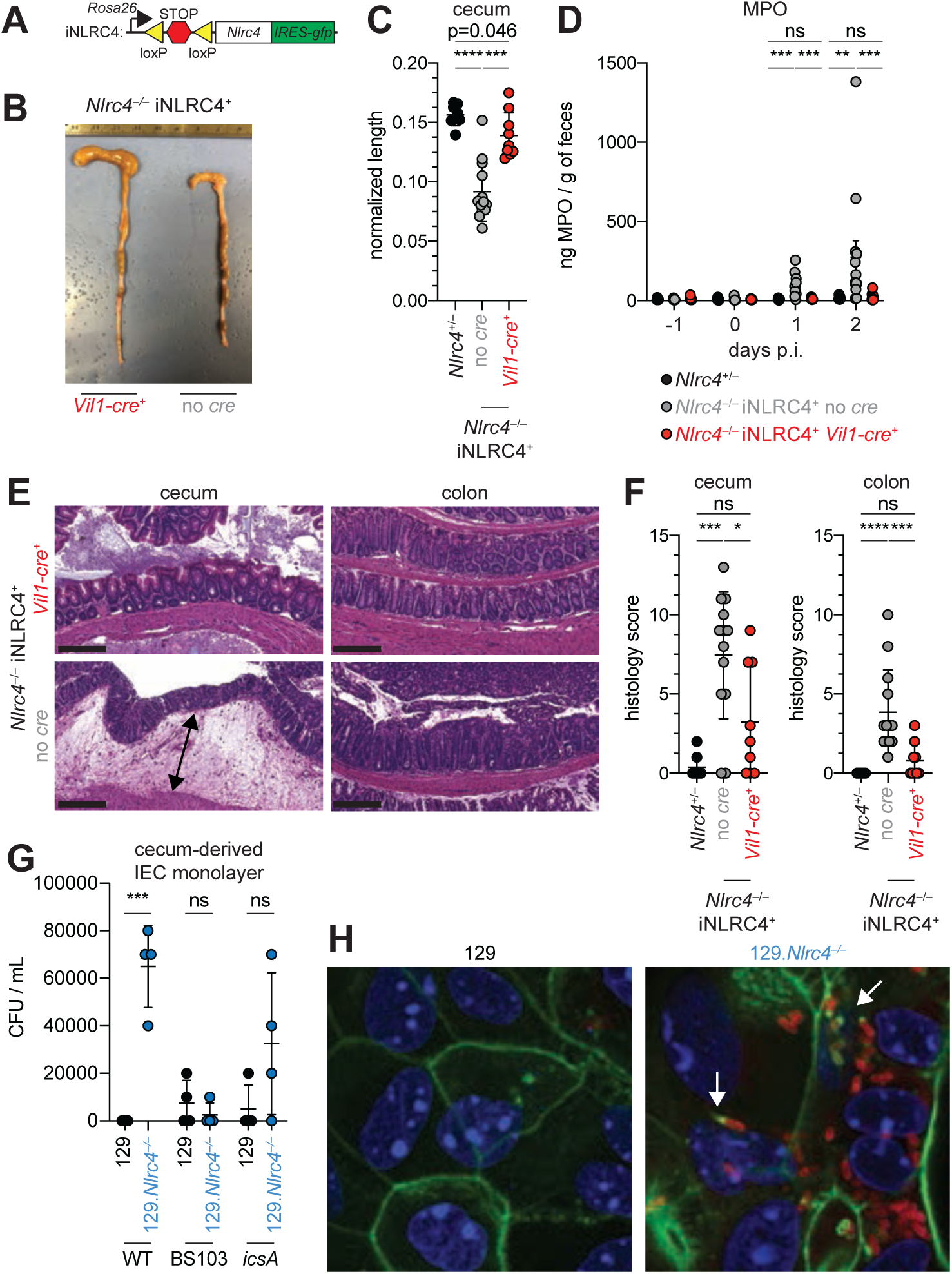
*Nlrc4* expression in IECs is sufficient to prevent shigellosis. (**A**) Schematic of the B6 *Rosa26* locus containing the iNLRC4 cassette, as described previously (Rauch et al., 2017). (**B–F**) Vil1-cre positive (+) or negative *Nlrc4*^*–/–*^ iNlrc4 littermates, or *Nlrc4*^+/–^ mice were orally infected with 5×10^7^ CFU of WT *Shigella* 24 hours after oral streptomycin treatment. Endpoint harvests were done 48 hours post-infection (p.i.). (**B**) Representative images of the cecum and colon dissected from iNLRC4 *Nlrc4*^*–/–*^ Vil1-cre positive or negative mice. (**C**) Quantification of cecum length reduction normalized to the weight of the animal prior to infection; cecum length (cm) / mouse weight (g). (**D**) MPO levels measured by ELISA of feces collected -1 through 2 days p.i. (**E**) Representative images of H&E stained cecum and colon tissue from infected mice. Scale bar, 200µm. (**F**) Blinded quantification of histology score (cumulative) for cecum and colon tissue. Data are representative of two independent experiments. Mean ± SD is shown in (**C,D,F**), Mann-Whitney test, \**P* < 0.05, \*\**P* < 0.01, \*\*\**P* < 0.001. (**C,D,F**) Each symbol represents one mouse. (**G**) *Shigella* (WT, BS103, or *icsA*) CFU from transwell culture of WT or 129.*Nlrc4*^*–/–*^ cecum-derived IEC monolayers. CFU was determined 8 hours p.i. Each symbol represents one infected monolayer. (**H**) Immunofluorescent staining of WT *Shigella* infected transwell cultures of WT or 129.*Nlrc4*^*–/–*^ cecum-derived IEC monolayers: green, fluorescent phalloidin (actin); red, anti-*Shigella* LPS, blue, DAPI (nucleic acid).

To further characterize the role of the NAIP–NLRC4 inflammasome during *Shigella* infection of IECs, we generated intestinal epithelial stem cell-derived organoids from the ceca of 129.WT and 129.*Nlrc4*^*–/–*^ mice, and established a transwell monolayer infection assay (see Methods). We were unable to recover CFUs from WT IEC monolayers infected with WT *Shigella* (**Figure 5G**) In contrast, 129.*Nlrc4*^*–/–*^ IEC monolayers supported replication of WT *Shigella*. The avirulent non-invasive BS103 strain was detected sporadically at low levels, independent of NAIP–NLRC4, while a strain lacking IcsA, a protein essential for *Shigella* actin tail formation and cell-to-cell spread (Bernardini et al., 1989; Goldberg and Theriot, 1995), colonized 129.*Nlrc4*^*–/–*^ IEC monolayers to a lesser extent than WT *Shigella*, consistent with loss of IcsA-mediated cell-to-cell spread. Immunostaining for *Shigella* in infected IEC organoid cultures also revealed intracellular replication and actin tail formation (detected by fluorescent phalloidin) exclusively in *Nlrc4*^*–/–*^ IEC monolayers infected with WT *Shigella* (**Figure 5H**). Thus, IEC organotypic infections faithfully recapitulate the NAIP–NLRC4-dependent differences in *Shigella* replication observed *in vivo*. We conclude that the protection mediated by NAIP–NLRC4 is cell-intrinsic and does not require cytokine signaling to additional immune cell populations, although such signaling may have additional effects *in vivo*. Importantly, these data further demonstrate that *Shigella* virulence factors are functional within mouse cells and can initiate invasion and actin-based motility in mouse IECs, as long as the NAIP–NLRC4 inflammasome is absent.

### IcsA-dependent cell-to-cell spread is required for pathogenesis

The *Shigella* IcsA protein is required for virulence in humans (Collins et al., 2008; Mani et al., 2016; Orr et al., 2005). To test if *icsA* is required for pathogenesis in mice, we infected 129.*Nlrc4*^−/−^ mice with isogenic WT, *icsA* mutant, or BS103 *Shigella* and monitored disease for eight days. Mice infected with WT *Shigella* exhibited weight loss (**Figure 6A**), diarrhea (**Figure 6B**), increases in fecal MPO (**Figure 6C**), and blood in their stool (**Figure 6D**). Signs of disease in WT-infected mice peaked between 2-3 days post-infection with weight loss, stool consistency, and MPO signal returning to baseline levels at approximately seven days post-infection, consistent with the disease progression and resolution of human shigellosis. Interestingly, 129.*Nlrc4*^−/−^ mice infected with *icsA* mutant *Shigella* did not experience weight loss, diarrhea, or fecal blood, and largely phenocopied mice infected with the non-invasive BS103 strain (**Figures 6A–D**). We did observe a slight but significant increase in fecal MPO levels at 1-3 days post-infection in these mice (**Figure 6B**). These results suggest that, as in humans (Coster et al., 1999; Kotloff et al., 1996), *icsA* mutants can provoke mild inflammation upon initial colonization of the intestinal epithelium, but that dissemination of bacteria among IECs is a critical driver of severe disease.

**Figure 6.**
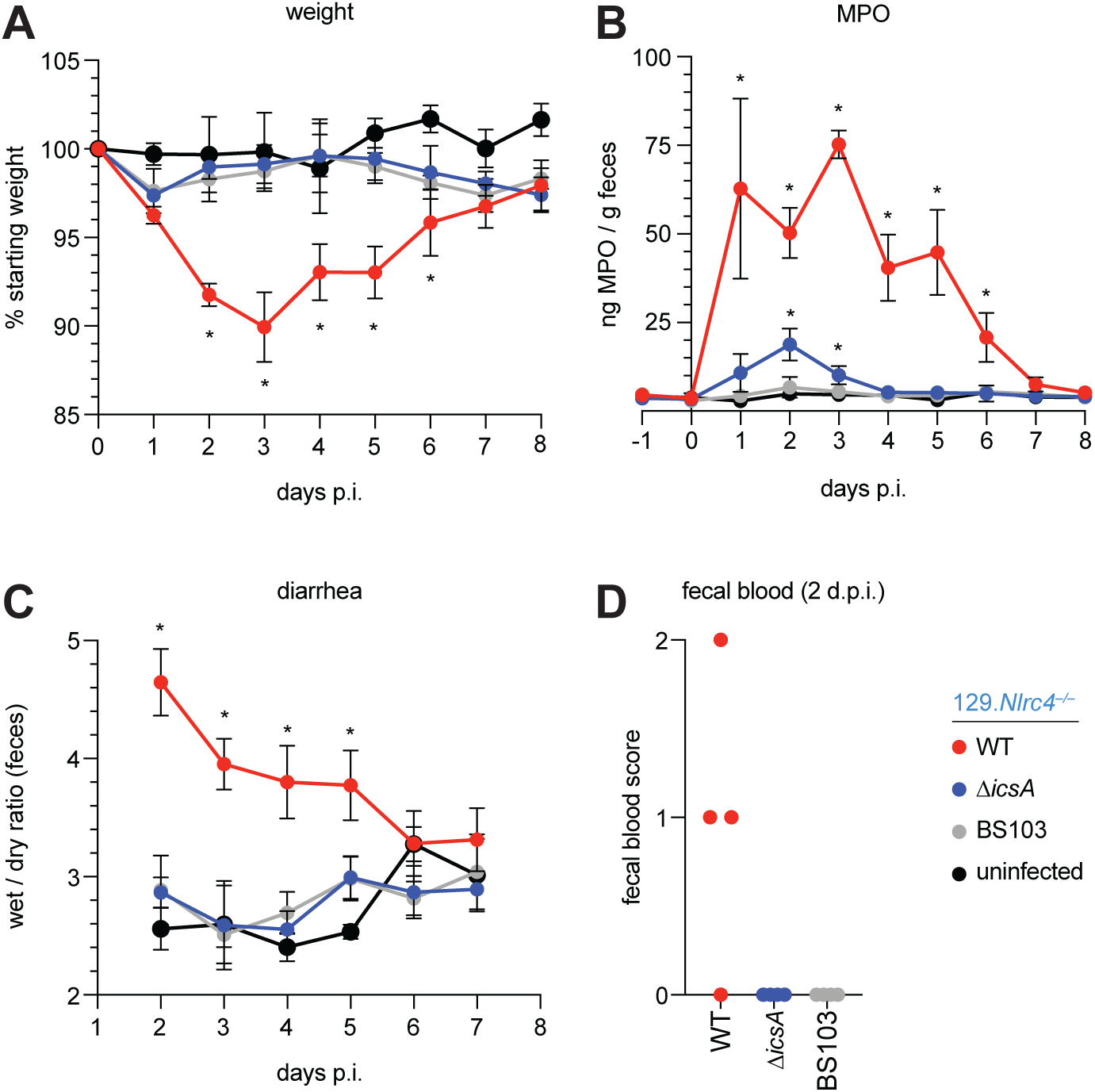
129.*Nlrc4*^*–/–*^ mice are resistant to attenuated *Shigella* strains. (**A–D**) 129.*Nlrc4*^*–/–*^ littermates were uninfected (black) or inoculated orally with 5×10^7^ CFU of WT (red), *icsA* mutant (blue), or BS103 (grey) *Shigella* 24 hours after oral streptomycin treatment and monitored for 8 days post-infection (p.i.). (**A**) Mouse weights. (**B**) MPO levels measured by ELISA from feces collected -1 through 8 days p.i. (**C**) Quantification of diarrhea comparing weight of feces before and after dehydration. A larger ratio indicates diarrhea. (**D**) Fecal blood scores from feces at two days post-infection (d.p.i.). 1 = occult blood, 2 = macroscopic blood. Each symbol represents feces from one mouse. (**A–C**) Each symbol represents the mean at a specific time point for four individual mice per infection condition. Data are representative of two independent experiments. Mean ± SEM is shown in (**A–C**) Mann-Whitney test, \**P* < 0.05. In (**A,B**) significance was determined by independently comparing to Day 0 and to BS103 + uninfected at the same day. In (**C**), significance was determined by comparing to BS103 + uninfected at the same day.

## Discussion

Here we demonstrate that the NAIP–NLRC4 inflammasome is a formidable species-specific barrier to *Shigella* invasion of the intestinal epithelium. *Shigella* infection of antibiotic pre-treated, NAIP–NLRC4-deficient mice recapitulates key features of human shigellosis, including bacterial invasion of and replication in IECs, severe inflammatory disease at relevant sites (e.g., colon, cecum), and bloody diarrhea. Thus, inflammasome-deficient mice provide the first physiologically relevant mouse model of bacillary dysentery, paving the way for genetic and mechanistic *in vivo* studies of the host factors underlying *Shigella* pathogenesis that have long been elusive.

A long-held belief is that *Shigella* exploits inflammasomes to induce pyroptosis. Pyroptotic cell death is presumed to allow bacteria to escape macrophages and invade the basolateral surface of polarized enterocytes (Ashida et al., 2014; Lamkanfi and Dixit, 2010; Schnupf and Sansonetti, 2019). Although we do not directly address this possibility, our experiments suggest that *Shigella* has instead evolved to inhibit the human NAIP–NLRC4 inflammasome to evade intestinal epithelial cell death. Indeed, we find that the NAIP–NLRC4 inflammasome plays a critical role in host defense by limiting *Shigella* replication and spread in IECs. Further supporting this notion, only *Nlrc4*^−/−^ but not WT IEC organoid monolayers are permissive to *Shigella* infection. *Salmonella* infected IECs are expelled from the intestinal epithelial barrier in an NLRC4-dependent manner (Rauch et al., 2017; Sellin et al., 2014). The same mechanism is also likely to occur during *Shigella* infection and suggests that epithelial inflammasomes coordinate the expulsion of infected IECs as a general defense strategy against enteric bacterial pathogens.

We only observe bloody diarrhea (dysentery) in *Nlrc4*^*–/–*^ mice generated on the 129 genetic background. 129 mice naturally harbor a null *Casp11* allele (Kayagaki et al., 2011). Although other genetic differences may contribute in part to the variation in susceptibility to *Shigella* between B6 and 129 NAIP–NLRC4-deficient mice, we speculate that both the NAIP–NLRC4 and the CASP11 inflammasomes mediate protection in IECs against *Shigella* invasion. However, the susceptibility of B6 NAIP–NLRC4-deficient (but CASP11^+^) mice, as well as the resistance of *Casp1/11*^−/−^ mice, suggest that these inflammasomes are not strictly redundant and that NAIP–NLRC4 alone is sufficient to confer resistance to shigellosis in mice. The *Shigella* effector OspC3 antagonizes the human LPS sensor CASP4 (Kobayashi et al., 2013), and IEC-expressed CASP4 provides protection against other human bacterial pathogens (Holly et al., 2020; Knodler et al., 2014), underscoring the importance of the LPS sensing pathway during human infection. Similarly, our finding that *Shigella* appears to suppress the human NAIP–NLRC4 inflammasome implies that evasion and/or antagonism of inflammasomes is a general virulence strategy during human infections.

There is currently no licensed *Shigella* vaccine, and very limited knowledge of what vaccine-induced immune responses would be desirable to elicit to mediate protection (Barry et al., 2013; Mani et al., 2016). Our new shigellosis model will finally allow the field to leverage the outstanding genetic tools and reagents in the mouse to address fundamental questions about the immune response to *Shigella*. Our finding that NAIP–NLRC4 inflammasome-deficient mice clear the attenuated *Shigella icsA* strain, derivatives of which are currently deployed in human vaccine trials (Collins et al., 2008; Coster et al., 1999; Ranallo et al., 2014), speaks to the readiness of our model to support testing and development of *Shigella* therapeutics. More broadly, our results also provide a striking example of how inflammasomes provide an important species-specific barrier against infection.

## Methods

### Cell culture

293T cells were cultured in DMEM supplemented with 10% FBS and 2mM L-glutamine. THP1 cells were cultured in RPMI supplemented with 10% FBS and 2mM L-glutamine. B6 primary BMMs were cultured in RPMI supplemented with 10% FBS, 5% mCSF, 100U/ml penicillin, 100mg/ml Streptomycin and 2mM L-glutamine. THP1 cells were a gift from Veit Hornung, and generated as previously described (Gaidt et al., 2017). Cells were grown in media without antibiotics for infection experiments.

### Bacterial strains

All experiments were conducted with the *S. flexneri* serovar 2a WT 2457T strain, or the WT-derived virulence plasmid-cured strain BS103 (Maurelli et al., 1984) or *icsA* mutant (Goldberg and Theriot, 1995; Makino et al., 1986). The *icsA* mutant strain was a gift from Marcia Goldberg. Natural streptomycin resistant strains of 2457T and BS103 were generated by plating cultured bacteria on tryptic soy broth (TSB) plates containing 0.01% Congo Red (CR) and increasing concentrations of streptomycin sulfate. Streptomycin-resistant strains were confirmed to grow indistinguishably from parental strains in TSB broth lacking antibiotics, indicating an absence of streptomycin-dependence.

### Toxins

Recombinant proteins for cytosolic delivery of *Shigella* MxiH were produced using the BD BaculoGOLD system for protein expression in insect cells. The MxiH coding sequence was subcloned into pAcSG2-6xHIS-LFn using the primers: PSMpr943 F (<u>BamHI</u>) 5’ - GAAAGG <u>GGATCC</u> ATG AGT GTT ACA GTA CCG GAT AAA GAT TGG ACT CTG - 3’ and PSMpr944 R (<u>NotI</u>) 5’ - GAAAGG <u>GCGGCCGC</u> TTA TCT GAA GTT TTG AAT AAT TGC AGC ATC AAC ATC C - 3’. The PA-6xHIS coding sequence was subcloned from pET22b-PA-6xHIS(Rauch et al., 2016) into pAcSG2 using the primers: PSMpr896 F (<u>XhoI</u>) 5’ - GAAAGG <u>CTCGAG</u> ATG GAA GTT AAA CAG GAG AAC CGG TTA TTA AAT GAA TC - 3’ and PSMpr897 R (NotI) 5’ - GAAAGG <u>GCGGCCGC</u> TCA GTG GTG GTG GTG GTG GTG T - 3’. Constructs were co-transfected with BestBac linearized baculovirus DNA (Expression Systems) into SF9 cells following the manufacturer’s protocol to generate infectious baculovirus. Primary virus was amplified in SF9 cells. Recombinant proteins were produced by infecting 2L of High Five cells with 1ml of amplified virus/L cells. Cells were harvested ∼60 hours after infection by centrifugation at 500x*g* for 15 minutes. Cell pellets were resuspended in lysis buffer (50mM Tris pH7.4, 150mM NaCl, 1% NP-40 with protease inhibitors) and lysed on ice using a dounce homogenizer. Samples were then clarified at 24,000x*g* for 30 minutes and supernatants were batch bound to 1ml nickel resin for 2 hours at 4°C. Samples were column purified by gravity. Resin was washed with 100ml of wash buffer (20mM Tris pH7.4, 400mM NaCl, 20mM imidizole). Sample was eluted with 1ml fractions of elution buffer (20mM Tris pH7.4, 150mM NaCl, 250mM imidizole). Peak elutions were pooled and buffer exchanged into 20mM Tris pH7.4.

### Infection of cells in culture

*S. flexneri* was grown at 37°C on tryptic soy agar plates containing 0.01% Congo red (CR), supplemented with 100μg/ml spectinomycin and 100 μg/ml carbenicillin for growth of the *icsA* strain. For infections, a single CR-positive colony was inoculated into 5ml TSB and grown shaking overnight at 37°C. Saturated cultures were back-diluted 1:100 in 5ml fresh TSB shaking for ∼2 hours at 37°C. THP1 and BMM cells were seeded at 100,000 cells per well of a Nunc F96 MicroWell white polystyrene plate. Bacteria were washed three times in cell culture media, then spinfected onto cells for 10 minutes at 500x*g*. Bacterial invasion was allowed to proceed for an additional 20 minutes at 37°C, followed by three washes in with cell culture media containing 25mg/ml gentamicin. Cells were then maintained in cell culture media containing 2.5mg/ml gentamicin with propidium iodide (Sigma, diluted 1:100 from stock) at 37°C for the duration of the assay (30 minutes to 4 hours). A MOI of 10 was used unless otherwise specified. For suppression assays, PMA-differentiated THP1 cells were infected as described above for 1 hour. Media was then replaced with cell culture media containing 2.5mg/ml gentamicin and propidium iodide (1:100) and either 10μg/ml PA only, PA with 1.0μg/ml LFn- MxiH, or 10μM nigericin. PI uptake was measured using a SpectraMax M2 plate reader, and 100% cell death was set by normalizing values of infected wells to cells lysed with 1% Triton X-100 after background subtraction based on media only controls.

### Establishment, propagation and infection of IECs

Primary intestinal epithelial stem cell-derived organoids from the cecum were isolated and maintained in culture as previously described(Miyoshi and Stappenbeck, 2013). Each transwell monolayer culture was established 1:1 from a confluent enteroid Matrigel (Corning, 356255) ‘dome.’ Enteroids were disassociated from Matrigel with 0.25% trypsin for 10 minutes, manually disrupted, resuspended in monolayer culture wash media (ADMEM/F12 supplemented with 20% FBS, 1% L-glutamine) and plated on polycarbonate transwells (Corning, 3413) that had been pre-coated for >1 hour at 37°C with 1:30 Matrigel:wash media. Monolayer cultures were differentiated for 12-14 days in complete monolayer culture media (monolayer culture wash media mixed 1:1 with LWRN-conditioned media supplemented with 10μM Y27632 (Stem Cell), and RANKL (BioLegend) in the absence of SB431542). Two days prior to infection, cells were cultured in antibiotic-free monolayer culture media. Monolayers were treated with 20μM EGTA for 15 minutes prior to *Shigella* infection (MOI=10). Bacterial invasion was allowed to proceed for 2 hours, followed by gentamicin washes as described above to both upper and lower compartments, then maintained in monolayer culture media containing 2.5mg/ml gentamicin with propidium iodide (Sigma, diluted 1:100 from stock) at 37°C for the duration of the assay (1 hour for IF and 8 hours for CFU determination). For IF, cells were washed in PBS, fixed in 4% paraformaldehyde for 15 minutes, permeabilized in 0.1% Triton X-100 for 15 minutes, blocked in PBS with 2% BSA, 0.1% Tween-20 for 1 hour. Primary antibodies were incubated overnight, followed by 1 hour stain with fluorophore-conjugated secondary antibodies and 10 minute staining with DAPI and fluorophore-conjugated phalloidin. Slides were analyzed on a Zeiss LSM710. Antibodies: anti-*Shigella* (Abcam, ab65282), 488 phalloidin (PHDG1-A, Cytoskeleton Inc.), Alexfluor conjugated secondary antibodies (Invitrogen). To determine bacterial replication in IECs, 8h post-infection monolayers were washed three times with PBS, lysed in 1% Triton X-100, and bacteria were plated for CFU determination.

### Reconstituted NAIP–NLRC4 inflammasome activity assays

To reconstitute inflammasome activity in 293T cells, constructs (100ng of each) producing human NAIP, NLRC4, CASP1 and IL-1β were co-transfected with constructs (200ng of each) producing *Shigella* MxiI, MxiH or empty vector (pcDNA3) using Lipofectamine 2000 (Invitrogen) following the manufacturer’s protocol and harvested 24 hours post-transfection. For experiments using recombinant proteins, fresh media containing 10μg/ml PA and 1.0μg/ml LFn-MxiH was added to cells for 3-4 hours. In all experiments, cells were lysed in RIPA buffer with protease inhibitor cocktail (Roche).

### Immunoblot and antibodies

Lysates were clarified by spinning at 16,100x*g* for 10 minutes at 4°C. Clarified lysates were denatured in SDS loading buffer. Samples were separated on NuPAGE Bis-Tris 4-12% gradient gels (ThermoFisher) following the manufacturer’s protocol. Proteins were transferred onto Immobilon-FL PVDF membranes at 375mA for 90 minutes and blocked with Odyssey blocking buffer (Li-Cor). Proteins were detected on a Li-Cor Odyssey Blot Imager using the following primary and secondary antibodies: anti-IL-1β (R&D systems, AF-201-NA), anti-GFP (Clontech, JL8), anti-TUBULIN (Sigma, clone TUB 2.1), Alexfluor-680 conjugated secondary antibodies (Invitrogen).

### Animal Procedures

All mice were maintained in a specific pathogen free colony until 1-2 weeks prior to infection, maintained under a 12 hour light-dark cycle (7am to 7pm), and given a standard chow diet (Harlan irradiated laboratory animal diet) ad libitum. Wild-type C57BL/6J and 129S1/SvImJ mice were originally obtained from the Jackson Laboratories. 129.*Nlrc4*^*–/–*^ animals were generated by targeting *Nlrc4* via CRISPR-Cas9 mutagenesis. CRISPR/Cas9 targeting was performed by pronuclear injection of Cas9 mRNA and sgRNA into fertilized zygotes, essentially as described previously(Wang et al., 2013). Founder mice were genotyped by PCR and sequencing using the primers: JLR035 F 5’ CAGGTCACAGAAG AAGACCTGAATG 3’ and JLR036 R 5’ CACCTGGACTCCTGGATTTGG 3’. Founders carrying mutations were bred one generation to wild-type mice to separate modified haplotypes. Homozygous lines were generated by interbreeding heterozygotes carrying matched haplotypes. B6.Δ*Naip* mice were generated as described previously (Rauch et al., 2016). B6.*Nlrc4*^*–/–*^ mice and iNLRC4 mice were generated as described previously(Rauch et al., 2017). iNLRC4 mice were crossed to the *Nlrc4*^*–/–*^ line and then further crossed to *Vil1*-cre (Jax strain 004586) transgenic lines on a *Nlrc4*^*–/–*^ background. Animals used in infection experiments were littermates or, if not possible, were co-housed upon weaning. In rare cases when mice were not co-housed upon weaning, mice were co-housed for at least one week prior to infection. Animals were transferred from a SPF colony to an ABSL2 facility at least one weeks prior to infection. All animal experiments complied with the regulatory standards of, and were approved by, the University of California, Berkeley Animal Care and Use Committee.

### *In vivo Shigella* infections

Mouse infections were performed in 6-16 week old mice. Initially, mice deprived of food and water for 4-6 hours were orally gavaged with 100µL of 250 mg/mL streptomycin sulfate dissolved in water (25 mg/mouse) and placed in a cage with fresh bedding. 24 hours later, mice again deprived of food and water for 4-6 hours were orally gavaged with 100µL of 5×10^8^ CFU (5×10^7^ CFUs per mouse) of log-phase, streptomycin resistant *Shigella flexneri* 2457T, BS103, or *icsA* mutant 2457T prepared as above and resuspended in PBS. Mouse weights and fecal pellets were recorded or collected daily from one day prior to infection to the day of euthanasia and harvest (usually 2 days post-infection) to assess the severity of disease and biomarkers of inflammation. Infection inputs were determined by serially diluting a fraction of the initial inoculum and plating on TSB plates containing 0.01% CR and 100µg/mL streptomycin.

### Fecal CFUs, fecal MPO ELISAs, wet/dry ratio, fecal occult blood

Fecal pellets were collected in 2mL tubes, suspended in 2% FBS in 1mL of PBS containing protease inhibitors, and homogenized. For CFU enumeration, serial dilutions were made in PBS and plated on TSB plates containing 0.01% CR and 100 µg/mL streptomycin sulfate. For MPO ELISAs, fecal homogenates were spun at 2,000*g* and supernatants were plated in triplicate on absorbent immunoassay 96-well plates. Recombinant mouse MPO standard, MPO capture antibody, and MPO sandwich antibody were purchased from R&D. Wet/dry ratios were determined by weighing fecal pellets before and after they had been dried in a fume hood. The presence or absence of fecal occult blood in fresh pellets was determined using a Hemoccult blood testing kit (Beckman Coulter).

### Histology

Mice were euthanized at two days post-infection by CO_2_ inhalation and cervical dislocation. Ceca and colons from mice were isolated, cut longitudinally, removed of lumenal contents, swiss-rolled, and fixed in methacarn followed by transfer to 70% ethanol. Samples were processed by routine histologic methods on an automated tissue processor (TissueTek, Sakura), embedded in paraffin, sectioned at 4µm thickness on a rotary microtome, and mounted on glass slides. Sections were stained with hematoxylin and eosin on an automated histostainer and coverslipped. Histopathological evaluation was performed by light microscopy (Olympus BX45, Olympus Corporation) at magnifications ranging from x20 to x600 by a board-certified veterinary pathologist (I.L.B.) who was blinded to the experimental groups at the time of evaluation. Representative images were generated as Tiff files from digitized histology slides scanned on a digital slide scanner (Leica Aperio AT2, Leica Biosystems). Images were taken using freely downloadable software (Image Scope, Leica Aperio, Leica Biosystems) and processed in Adobe Photoshop. Photo processing was confined to global adjustments of image size, white balance, contrast, brightness, sharpness, or correction of lens distortion and did not alter the interpretation of the image. Sample preparation, imaging, and histology scoring was conducted by the Unit for Laboratory Animal Medicine at the University of Michigan.

### Intestinal CFU determination

To enumerate whole tissue intestinal CFU, ceca and colons from mice were isolated, cut longitudinally and removed of lumenal contents, placed in culture tubes containing 400µg/mL gentamicin antibiotic in PBS, vortexed, and incubated in this solution for 1-2 hours. Organs were washed 5 times in PBS to dilute the gentamicin, homogenized in 1mL of PBS, serially diluted, and plated on TSB agar plates containing 0.01% CR and 100µg/mL streptomycin. To enumerate intracellular CFU from the intestinal epithelial cell fraction of the cecum and colon, organs prepared as above were incubated in RPMI with 5% FBS, 2mM L-glutamine, and 400µg/ml of gentamicin for 1-2 hours. Tissues were then washed 5 times in PBS, cut into 1cm pieces, placed in 15mL of stripping solution (HBSS, 10mM HEPES, 1mM DTT, 2.6mM EDTA), and incubated at 37°C for 25 minutes with gentle agitation. Supernatants were passed through a 100 µm filter and the remaining pieces of tissue were shaken in a 50mL conical with 10 mL of PBS and passed again through the 100µm filter. This enriched epithelial cell fraction was incubated in 50µg/mL gentamicin for 25 minutes on ice, spun at 300x*g* at 4°C for 8 minutes, and washed twice by aspirating the supernatant, resuspending in PBS, and spinning at 300x*g* at 4°C for 5 minutes. After the first wash, a fraction of cells were set aside to determine the cell count. After the second wash, the pellet was resuspended and lysed in 1mL of 1% Triton X-100. Serial dilutions were made from this solution and plated on TSB agar plates containing 0.01% CR and 100µg/ml streptomycin and CR+ positive colonies were counted following overnight incubation at 37°C.

## Acknowledgements

We thank M. Goldberg for advice and for sharing the *icsA* mutant *Shigella* strain. We are grateful to G. Barton and H. Darwin for comments on the manuscript, and members of the Vance and Barton Labs for discussions. Funding: R.E.V. is an HHMI Investigator and is supported by NIH AI075039 and AI063302; P.S.M. is supported by a Jane Coffin Childs Memorial Fund postdoctoral fellowship. J.L.R. is an Irving H. Wiesenfeld CEND Fellow; E.A.T. is supported by the UC Berkeley Department of Molecular and Cell Biology NIH Training Grant 5T32GM007232-42; C.F.L. is a Brit d’Arbeloff MGH Research Scholar and supported by NIH AI064285 and NIH AI128743; I.R. is supported by the Medical Research Foundation MRF2012.

## Competing interests

R.E.V. has a financial relationship with Aduro BioTech and Ventus Therapeutics and both he and the companies may benefit from the commercialization of the results of this research.

## Author contributions

P.S.M, J.L.R., and R.E.V. conceived the study, designed the experiments, and wrote the original manuscript; P.S.M and J.L.R. performed the majority of the experiments with contributions from E.A.T and R.A.C.; I.R., L.G., and C.F.L. provided resources; I.R. and C.F.L. contributed to methodology and supervision; P.S.M, J.L.R., R.E.V., I.R., L.G., and C.F.L. edited and reviewed the manuscript.

---

**Figure S1.**
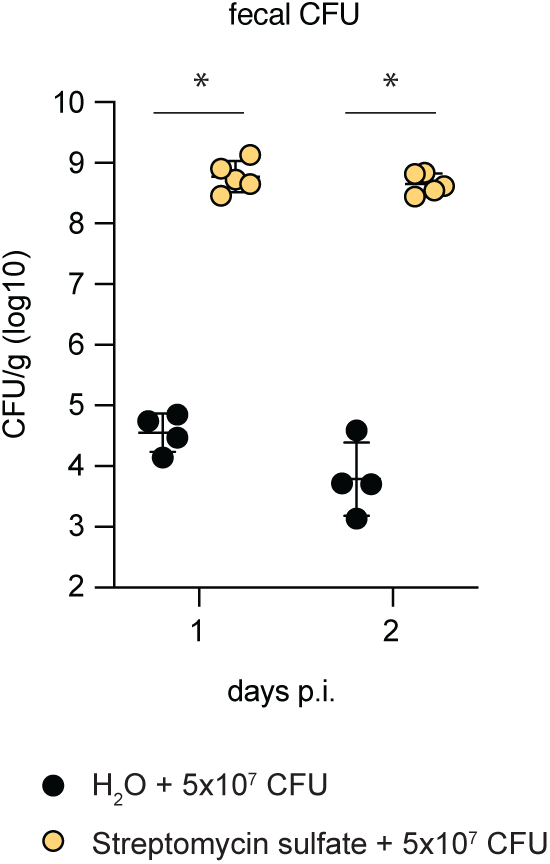
Antibiotic pre-treatment followed by oral route *Shigella* infection permits substantial lumenal colonization. CFU determination per gram (g) feces of B6 WT mice that were treated orally with 25mg streptomycin sulfate (yellow) or water (black) and orally challenged the next day with 5×10^7^ CFU of WT *Shigella*. Feces were collected 1 and 2 days post-infection (p.i.). Data are representative of three experiments. Each symbol represents one mouse. Mann-Whitney test, \**P* < 0.05.

**Figure S2.**
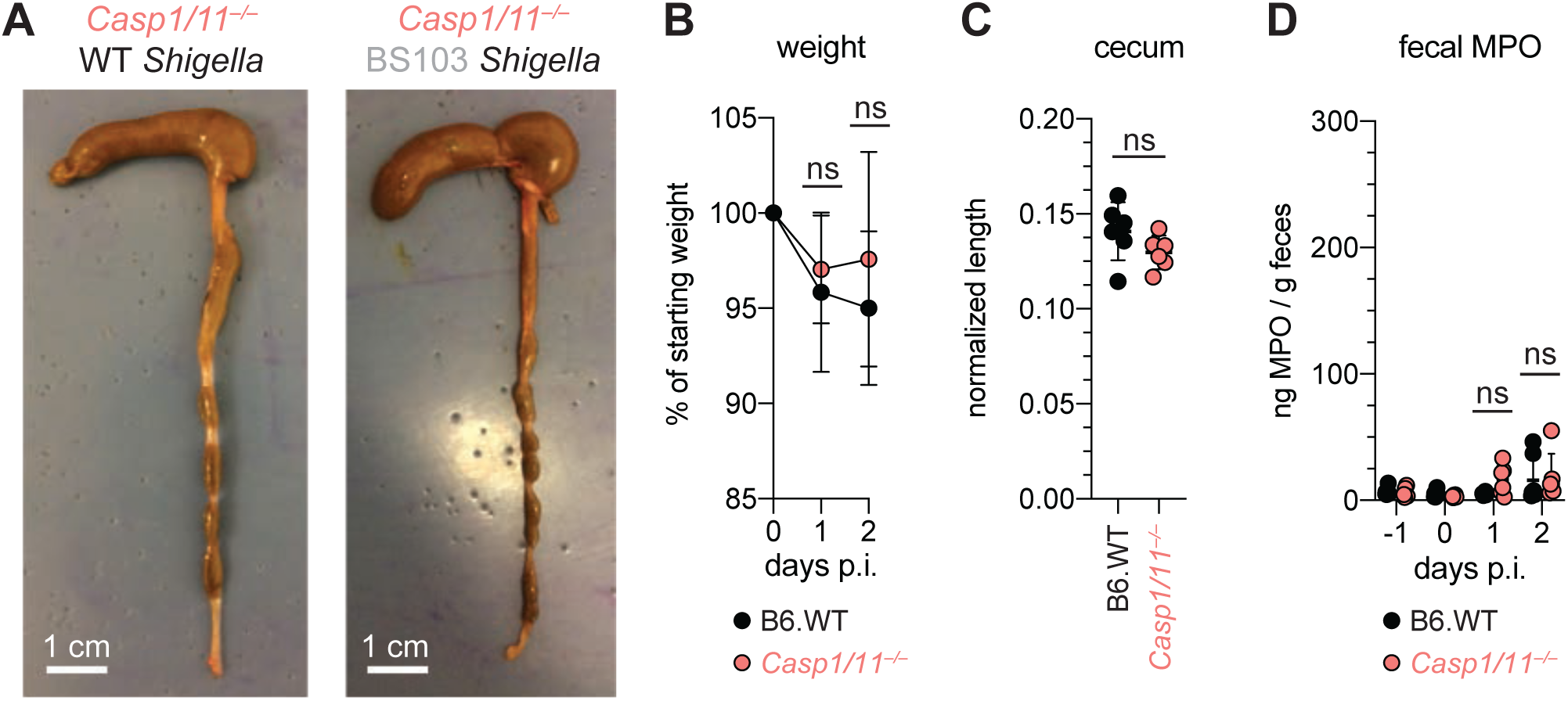
B6.*Casp1/11*^*–/–*^ mice are resistant to oral *Shigella* challenge. (**A–D**) B6.WT (black) and B6.*Casp1/11*^***–/–***^ (peach) mice were treated orally with 25mg streptomycin sulfate and were orally challenged the next day with 5×10^7^ CFU of either WT or BS103 (avirulent) *Shigella*. (**A**) Representative images of the cecum and colon dissected at 2 days post-infection (p.i.). (**B**) Mouse weights. (**C**) Quantification of cecum and colon lengths. Values were normalized to mouse weight prior to infection; cecum length (cm) / mouse weight. (**D**) MPO levels measured by ELISA from feces of B6.WT and B6.*Casp1/11*^***–/–***^ mice collected -1 through 2 days p.i. Each symbol represents one mouse. Data are representative of three independent experiments. Mean ± SD is shown in (**B,C,D**), Mann-Whitney test, \**P* < 0.05.

**Figure S3.**
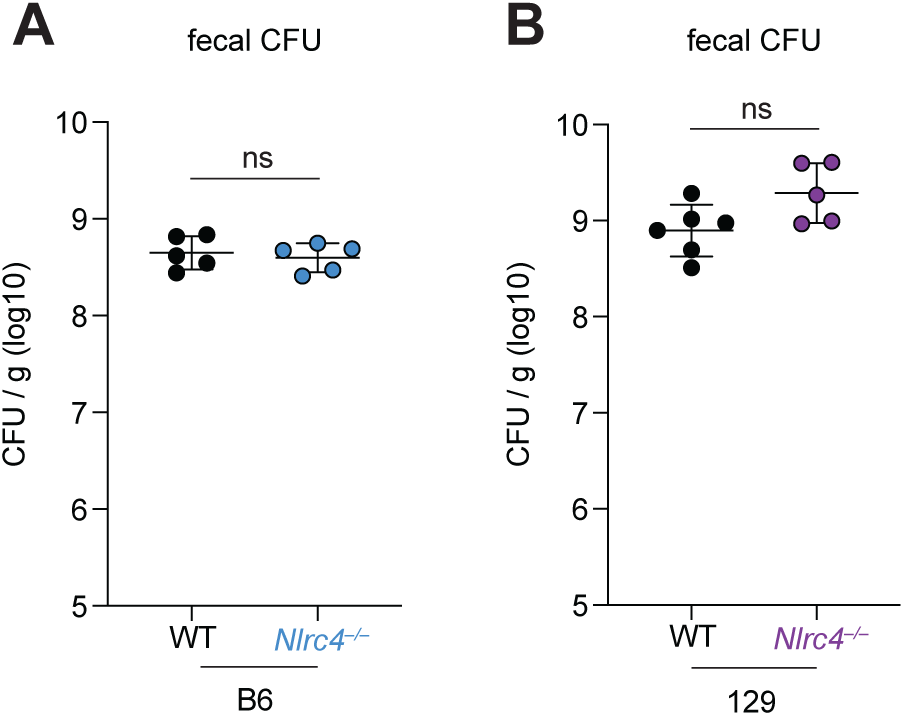
Lumenal colonization by *Shigella* is similar between WT and NAIP–NLRC4-deficient mice. (**A**) CFU determination per gram (g) feces from B6 and B6.*Nlrc4*^*–/–*^ *Shigella*-infected mice 2 days post-infection. (**B**) CFU determination per gram (g) feces from 129 and 129.*Nlrc4*^*–/–*^ *Shigella*-infected mice 2 days post-infection. Each symbol represents one mouse. Mann-Whitney test, \**P* < 0.05.

**Figure S4.**
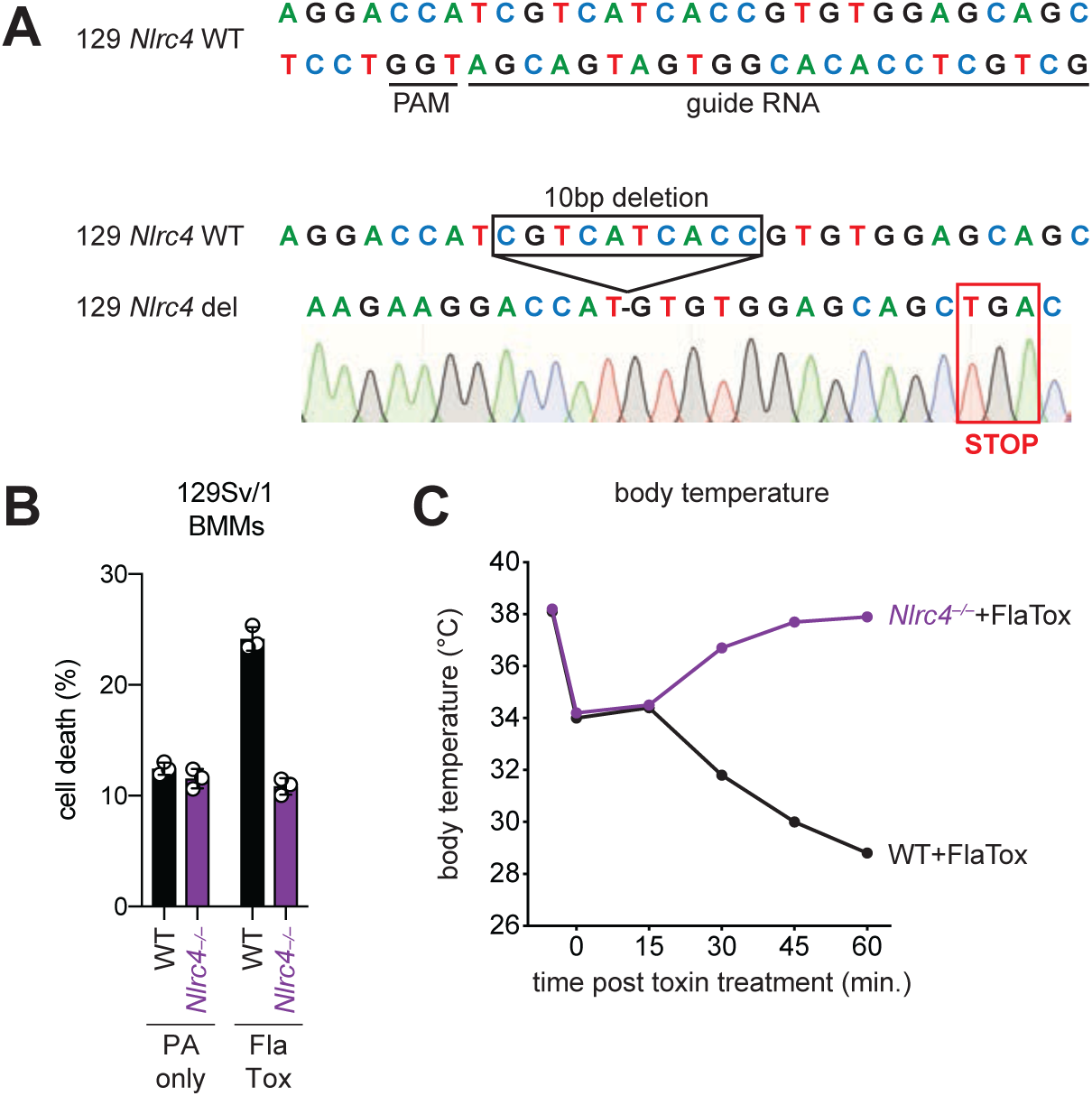
Construction and functional characterization of *Nlrc4* knockout mice on the 129S1/SvImJ genetic background. (**A**) The targeted wildtype *Nlrc4* sequence (chromosome 17, NC_000083.6, exon 5) aligned to the *Nlrc4* guide RNA. The protospacer adjacent motif (PAM) is indicated. Below is a schematic of the Sanger sequencing verified product of CRISPR/Cas9-editing (129 *Nlrc4* del), which results in a 10 base pair deletion and an in-frame early TGA stop codon in exon 5 of *Nlrc4*. (**B**) Quantification of cell death in 129 WT or *Nlrc4*^*–/–*^ bone marrow derived macrophages (BMMs) treated with 10µg/mL PA alone or PA + 10µg/mL LFn-FlaA (LFn fused to *Legionella pneumophila* flagellin, “FlaTox”). Cell death was measured 30 minutes post-infection by propidium iodide uptake and reported as percent death relative to 100% killing by treatment with Triton X-100. (**C**) WT or 129.*Nlrc4*^*–/–*^ mice were injected intravenously with 0.2µg/g body weight PA + 0.1µg/g body weight LFn-FlaA and body temperature was monitored for the indicated times (minutes) post-treatment. The initial temperature decrease in all mice is due to isoflurane treatment.

## Notes

### Competing Interest Statement

REV has a financial relationship with Aduro Biotech and both he and the company may benefit from commercialization of the results of this research

